# Distinct tooth regeneration systems deploy a conserved battery of genes

**DOI:** 10.1101/2020.09.21.305466

**Authors:** Tyler A. Square, Shivani Sundaram, Emma J. Mackey, Craig T. Miller

**Affiliations:** Department of Molecular & Cell Biology, University of California, Berkeley, United States

**Keywords:** Successional Dental Lamina, Tooth Regeneration, Odontode, Epithelial Appendage

## Abstract

**Background:** Vertebrate teeth exhibit a wide range of regenerative systems. Many species, including most mammals, reptiles, and amphibians, form replacement teeth at a histologically distinct location called the successional dental lamina, while other species do not employ such a system. Notably, a ‘lamina-less’ tooth replacement condition is found in a paraphyletic array of ray-finned fishes, such as stickleback, trout, cod, medaka, and bichir. Furthermore, the position, renewal potential, and latency times appear to vary drastically across different vertebrate tooth regeneration systems. The progenitor cells underlying tooth regeneration thus present highly divergent arrangements and potentials. Given the spectrum of regeneration systems present in vertebrates, it is unclear if morphologically divergent tooth regeneration systems deploy an overlapping battery of genes in their naïve dental tissues.

**Results:** In the present work, we aimed to determine whether or not tooth progenitor epithelia could be composed of a conserved cell type between vertebrate dentitions with divergent regeneration systems. To address this question, we compared the tooth regeneration processes in two ray-finned fishes: zebrafish (*Danio rerio*) and threespine stickleback (*Gasterosteus aculeatus*). These two teleost species diverged approximately 250 million years ago, and demonstrate some stark differences in dental morphology and regeneration. Here we find that the successional dental lamina in zebrafish sharply upregulates Wnt signaling and *Lef1* expression during early morphogenesis stages of tooth development. Additionally, the naïve zebrafish successional dental lamina expresses a battery of nine genes (*Bmpr1a, Bmp6, CD34, Gli1, Igfbp5a, Lgr4, Lgr6, Nfatc1*, and *Pitx2*). We also find that, despite the absence of a histologically distinct successional dental lamina in stickleback tooth fields, new tooth germs also sharply upregulate Wnt signaling and *Lef1* expression, and additionally express the same battery of nine genes in the basalmost endodermal cell layer from which replacement tooth epithelia arise. Thus, two fish systems that either have an organized successional dental lamina (zebrafish) or lack a morphologically distinct successional dental lamina (sticklebacks) deploy similar genetic programs during tooth regeneration.

**Conclusions:** We propose that the expression domains described here delineate a highly conserved “successional dental epithelium” (SDE). Furthermore, a set of orthologous genes is known to mark hair follicle epithelial stem cells in mice, suggesting that regenerative systems in other epithelial appendages may utilize a related epithelial progenitor cell type, despite the highly derived nature of the resulting functional organs.

## Background

Vertebrate teeth arose shortly after vertebrates themselves and have since greatly diversified. Vertebrate “odontodes”, which include teeth and denticles, are defined by a shared basic morphology: a dentine core derived from mesenchymal cells, and an enamel or enameloid tip sheath contributed (at least in part) by epithelial cells [1–4]. The first clear evidence of dentine and enameloid-bearing structures arose at least ∼420 million years ago in fossils of placoderms, though an even earlier origin in conodonts has been debated [5–8]. Support for the homology of odontodes comes from developmental studies that document similar histogenesis, morphological development, and shared gene expression patterns between teeth and other odontodes across large phylogenetic distances within jawed vertebrates [9–12]. Despite these deeply-conserved, fundamental aspects of tooth and odontode development, teeth have radiated vastly in their arrangement [13], shape [14], size [15], placement within the body plan [16], and the regeneration system they employ [17,18]. Thus, some aspects of tooth development are staunchly conserved, while others are wildly plastic in evolution. This variation in the conservation of different aspects of tooth development presents an interesting case study in the evolution of development: what genetic signatures, if any, are common to disparate dental morphologies? Here, we assess two divergent tooth regeneration strategies found in two different ray-finned fishes: zebrafish (*Danio rerio*) and threespine stickleback (*Gasterosteus aculeatus*).

The morphogenesis of individual tooth organs has been well-studied across living jawed vertebrates [9,19]. Teeth form like other epithelial appendages, demonstrating an intricate epithelial-mesenchymal interaction and coordinated cell motions during their early development (Fig. 1) [20]. Primary teeth first appear as a placode (thickened epithelium) underlain by a mesenchymal condensation, followed by invagination, cap formation, and eventually deposition of dentine and enamel and/or enameloid, which are both composed of highly-concentrated hydroxyapatite (a hard, crystalline calcium phosphate mineral that gives bones their characteristic hardness) [3,21]. Some vertebrates, such as mice, do not replace entire teeth during their lives. By contrast, in most known extant vertebrate dental systems, once a tooth position is established, tooth replacement via regeneration can occur anywhere between one (e.g. most mammalian teeth) to potentially one hundred times or more (e.g. some sharks) at a given tooth position [22–27]. In many vertebrates with tooth replacement, this process initiates at a site called the successional dental lamina (SDL). The SDL can be identified histologically as a deep extension of epithelium connected directly to the outer dental epithelium of a predecessor tooth [9,28]. When present, during tooth replacement the SDL first thickens, similar to the placode stage of primary teeth [9]. Subsequently, this thickened epithelium undergoes the stereotypical tooth differentiation process outlined above (depicted in Fig. 1), and at all stages is tightly associated with underlying mesenchyme. The SDL was recently shown to be a key tissue in tooth regeneration: mouse molars, which do not naturally undergo regeneration, can be prompted to do so by stabilizing Wnt signaling in the rudimentary mouse SDL [29]. In most cases, a replacement tooth germ eventually prompts osteoclast activity to enzymatically dissociate the base of the predecessor tooth and remodel the underlying bone of attachment (if present) either before or during early eruption [30–34]. This osteoclast activity facilitates the shedding of the predecessor and makes space for the replacement tooth to finish mineralization and eruption, thus completing the regenerative process.

**Fig. 1.**
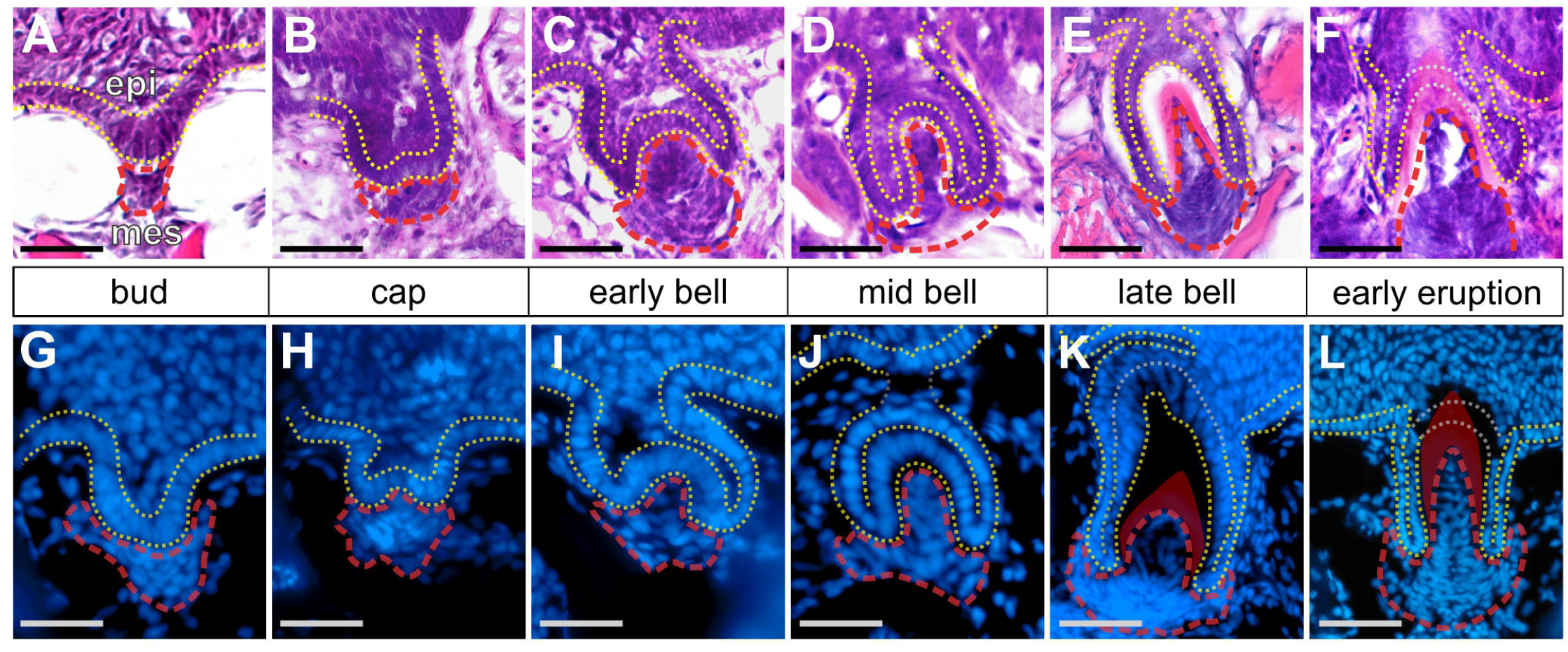
Stickleback replacement tooth morphogenesis. All images show sagittal or transverse sections from 20+ mm standard length sticklebacks. The basalmost layer of epithelium is outlined with yellow dashed lines in all panels, the mesenchyme is outlined in red. During late-bell stages (E, K), the epithelium towards the apex of the forming tooth becomes thickened, no longer presents a clear bilayer, and will eventually be punctured during tooth eruption (gray outlines). **A-F** Staging series of hematoxylin and eosin-stained transverse sections through stickleback replacement tooth germs at the stages indicated. Epithelium (“epi”) and mesenchyme (“mes”) are labeled in A. **G-L** Staging series of DAPI-stained sagittal sections through stickleback replacement tooth germs at the stages indicated. Gray portions of dotted epithelium lines in J indicate the epithelium/tooth connection that was present on an adjacent section. Bone matrix is false-colored red in K and L. Scale bars = 20 μm.

While many vertebrate tooth regeneration systems utilize an SDL, the arrangement of primary and replacement teeth is variable, both in modern species [9,12,17,28,35–41], as well as stem group jawed vertebrates [42,43]. In some species (e.g. sharks), teeth are replaced in families from permanent, conspicuous regions in the oral jaw [12]. By contrast, in other species, teeth are replaced from a relatively inconspicuous SDL that forms transiently (as in zebrafish) [39]. Some fishes such as bichir and trout present an even more abbreviated process, wherein the outer dental epithelium of the predecessor tooth directly relocalizes to a deeper region on the lingual side of the tooth, undergoes thickening, and eventually tooth differentiation, effectively ‘skipping’ the SDL phase [18,44]. Cichlid oral teeth grossly undergo the same process, though interestingly their oral replacement teeth are derived from the opposite (labial) side of the predecessor [10,45]. The patterning of tooth replacement can even vary throughout ontogeny of the same species: medaka fish initially have disorganized pharyngeal dentitions without visible tooth families, but later in life the teeth organize into highly-patterned pharyngeal tooth plates with clear tooth families arranged in rows [35,46]. Interestingly, despite these clear tooth families, no discrete dental lamina was observed in medaka [35]. Together, these data show that much flexibility exists in how progenitors can be arranged across dental regeneration systems [47].

Despite high variation in the size, shape, and position of vertebrate teeth, gene expression and functional studies on tooth differentiation across vertebrates has revealed deeply-conserved molecular processes at play [10,47]. However, given the phylogenetic lability and morphological disparities of tooth regeneration strategies between some groups [42], it is unclear whether or not this process is homologous and uses shared cell types across vertebrates. Recent evidence suggests that not just teeth, but also other epithelial appendages (EAs) such as scales, feathers, and hair share some aspects of development, also raising the possibility of far-reaching EA homology among these organs [20,48–55].

To further elucidate the developmental program underlying tooth regeneration, we compared the histology and gene expression patterns associated with the regenerative process in two teleost fishes: zebrafish and threespine stickleback. These two species share a common ancestor that lived approximately 250 million years ago [56] and appear to demonstrate some stark differences in their dentitions. Sticklebacks are well documented as demonstrating significant differences in tooth number between populations, particularly between high-toothed freshwater populations and low-toothed marine populations [57–61]. These tooth number differences have a strong genetic basis, contributing to a stark difference in adult tooth number between certain populations, which is especially pronounced on the ventral pharyngeal jaw [59,60]. We find that, unlike other vertebrates like zebrafish, the threespine stickleback neither exhibits an SDL nor regenerates teeth via the relocalization of the deep outer dental epithelium of a predecessor tooth (as in salmonid and bichir oral teeth). Furthermore, stickleback teeth appear to occasionally diverge from the common one-for-one mode of tooth replacement: some tooth germs appear to prompt osteoclast activity and shedding of more than one erupted tooth in concert. Despite these differences in modes of tooth regeneration, we find a suite of genes expressed in common between naïve dental epithelial cells in both fish species, suggesting that a conserved epithelial progenitor cell type underlies tooth regeneration, here referred to as the “successional dental epithelium” (SDE). We propose the use of this term to encompass not just the progenitor cells found in the histologically distinct successional dental lamina (SDL) of animals such as zebrafish, sharks, and tetrapods (where present), but also these less pronounced epithelial regions in fishes like sticklebacks, which still undergo constant tooth regeneration across their tooth fields. Intriguingly, the genes we find localized to the SDE of both zebrafish and sticklebacks also mirror a cassette of gene expression that is well documented and functionally relevant in hair follicle epithelial stem cells (in the “bulge”) [62,63], suggesting that such a genetic module may be common across not just teeth, but also other vertebrate epithelial appendages.

## Results

### 1. A brief review and comparison of the gross morphologies of stickleback and zebrafish dentitions

#### a. Tooth field position and arrangement

Zebrafish and stickleback teeth are grossly similar in tooth shape and structure (Fig. 2A and B); both species’ teeth are unicuspid and mostly conical, with a slight bend or hook towards the posterior side. However, these two species differ drastically in tooth arrangement within the body plan. Zebrafish, like other members of Cypriniformes, lack oral teeth, and possess only a single (paired) tooth field found ventrally on the last branchial arch (on ceratobranchial 5) in the pharynx (See Fig. 2A, referred to here as the ventral tooth plate or VTP, also called the lower pharyngeal jaw or LPJ in some species). Sticklebacks, like most vertebrates, have oral teeth, which are located on their dorsal premaxilla and ventral dentary bones. In the pharynx, sticklebacks also have a single paired VTP on ceratobranchial 5 (See Fig. 2B), as well as two paired dorsal tooth plates (DTPs) [64].

**Fig. 2.**
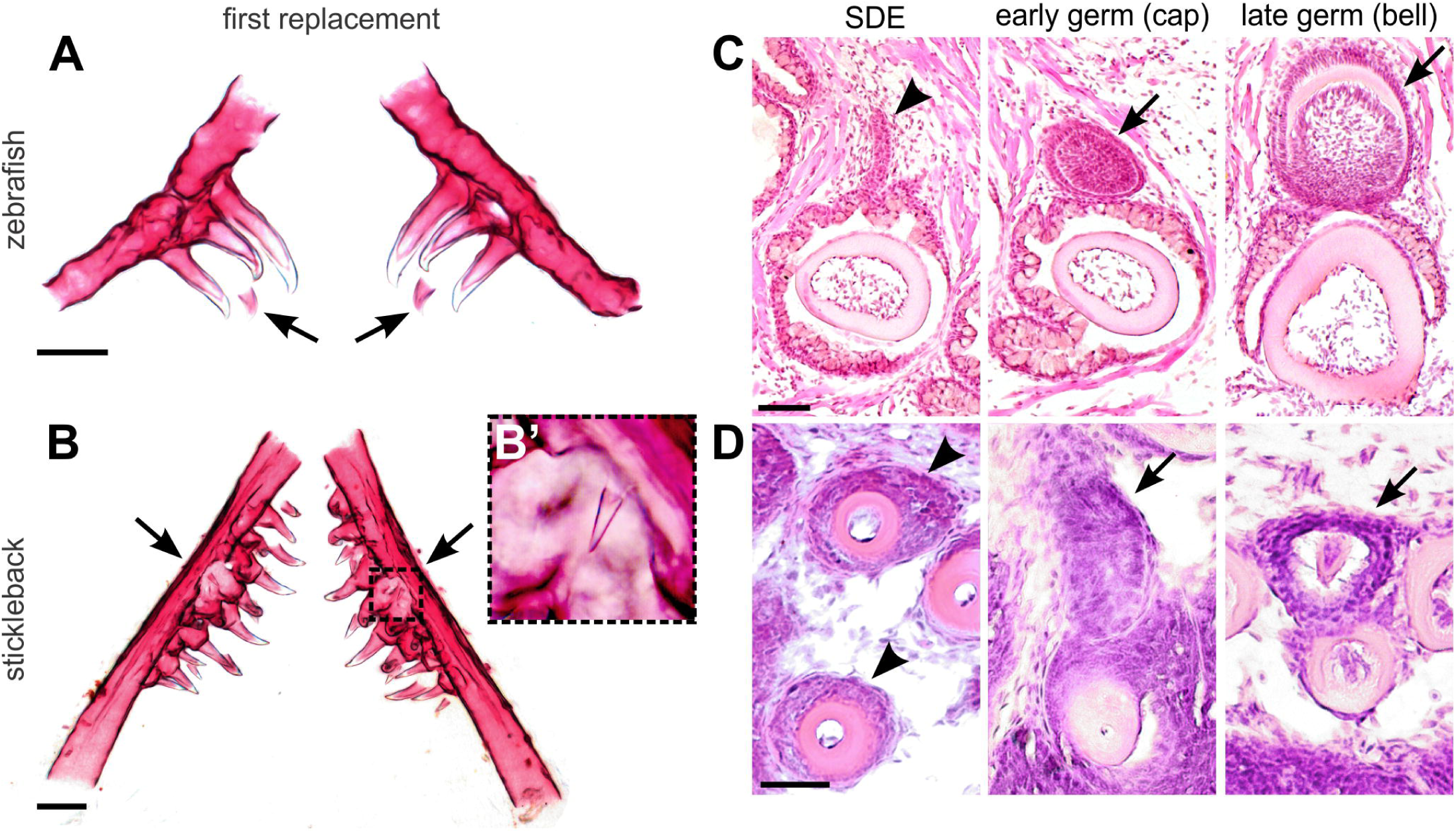
A comparison of zebrafish and stickleback tooth replacement. **A, B** Zebrafish and stickleback first tooth replacement events, respectively. Alizarin stained and dissected ceratobranchial 5 elements and their teeth from both species at 4 dpf zebrafish (A) and 30 dpf stickleback (B). Anterior to top. Arrows indicate first replacement tooth germs (bilaterally paired). Note that the zebrafish germs form on the ventral/medial side of the predecessor pioneer tooth (which in this case is the middle of 3 the ankylosed teeth on each side, tooth 4v1). Inset box in B magnified in B’ shows first replacement tooth. **C, D** Replacement histology on the coronal (D) and oblique coronal (C) axes (see methods). Arrowheads mark the successional dental epithelium (SDE). In zebrafish, this tissue takes the form of a true successional dental lamina (SDL). In sticklebacks, there is only a subtle, non-elongated epithelial thickening that surrounds the tooth shaft like a collar, immediately beneath the rest of the oral epithelia (see Fig. S1 for sagittal view). Arrows mark tooth germs at the stages indicated. Scale bars in A and B = 100 μm; C and D = 20 μm.

#### b. Tooth number variation

Sticklebacks and zebrafish also have major differences in tooth number. In lab-reared zebrafish, each VTP typically hosts 11 tooth positions (per side, so 22 total) (Van der Heyden and Huysseune 2000). As in other cyprinids, this number is known to vary occasionally in zebrafish, occasionally observed as missing or bearing an extra tooth position [65]. Since replacement tooth mineral deposition occurs prior to the shedding of a predecessor tooth during the tooth regeneration process, an adult zebrafish typically hosts 15-17 mineralized teeth per VTP at any given point in time (i.e. there are typically 4-6 late bell stage replacement tooth germs per VTP that have not yet dislodged their predecessor). Given this relatively consistent dental arcade, zebrafish tooth positions have been named based on their location [66].

Threespine sticklebacks, on the other hand, can have up to ∼200-300+ total teeth once fully grown, thus displaying both an order of magnitude more teeth than zebrafish, as well as higher within-species variation in adult tooth number. Tooth number in stickleback has a strong genetic basis, and varies significantly by population, with freshwater localities generally possessing more teeth than their marine counterparts [57,59–61].

#### c. Early replacement tooth events

Sticklebacks and zebrafish both appear to replace their earliest primary teeth in a similar order, with those teeth that differentiate first also being replaced first [64,66]. Interestingly, the tooth germs responsible for these early replacement events are essentially locationally opposite between the zebrafish and stickleback VTPs (arrows in Fig. 2A and B): zebrafish replacement tooth germs form on the medial/ventral side of each erupted tooth, towards the midline [66], while the first stickleback replacement tooth germs form on the lateral edge of the tooth plate, opposite the midline [64]. In zebrafish, this stereotypical polarized replacement direction (Fig. 2A) remains into adult stages. Conversely, as they mature, stickleback pharyngeal tooth fields do not appear to strictly adhere to the ‘lateral to medial’ replacement direction found in their earliest replacement events, which we detail below.

#### d. Replacement tooth histology

The histology of zebrafish tooth regeneration has been described previously in detail [39,66]. Zebrafish tooth regeneration initiates from an SDL. In zebrafish, this region forms transiently, as an outpocketing on the ventro-medial side of each tooth, first visible as a thin projection of naïve epithelial cells stemming from the outer dental epithelium of a predecessor tooth (arrowhead in Fig. 2C) [39], protruding from the mineralized tooth and its associated epithelia deep in the tooth field.

Stickleback pharyngeal replacement teeth arise from a more superficial region of the dentition than replacement teeth in zebrafish. After morphogenesis (Fig. 1), once ankylosis and eruption are complete, stickleback teeth do not retain a histologically distinct inner and outer dental epithelium (Fig. S1). Instead, mature stickleback pharyngeal teeth retain a shallow “collar” of epithelial cells surrounding the site of eruption (arrowheads in Fig. 2D and S1) that does not appear to be arranged in a clear bilayer. This collar of cells is contiguous with the basalmost epithelial sheet overlying the tooth field, juxtaposed to the pharyngeal epithelium (towards the lumen) and the basement membrane overlying mesenchymal cells (away from the lumen). It is from this collar of cells that replacement teeth appear to be derived (arrows in Fig. 2D). Thus, a histologically distinct SDL is not present during the regeneration of pharyngeal teeth in sticklebacks, a condition also reported for the oral teeth of bichir [18], cod [67], and trout [44], as well as pharyngeal teeth in medaka [35].

#### e. Stickleback pharyngeal teeth do not exhibit a tightly regulated 1-for-1 replacement scheme

Unlike zebrafish, stickleback tooth fields do not maintain any obvious arrangement of individual tooth positions across the entirety of each tooth field into adulthood, even within single populations. This observation initially led us to hypothesize that the stickleback tooth replacement process may introduce some spatial stochasticity of tooth position and spacing, similar to the condition in cod oral teeth, where their adult arrangement appears “haphazard” [67]. We thus sought to test the hypothesis that stickleback tooth replacement can diverge from a strict 1-for-1 tooth replacement system. We did this by analyzing H&E-stained serial sections cut in the sagittal, transverse, and coronal planes through pharyngeal tooth fields from subadult and adult fish (25-40 mm standard length; see Fig. S1 for section plane orientation). Using these preparations, we identified every putative pharyngeal and oral replacement tooth germ that was in a mid-to late-bell stage (the stages that coincide with bone remodeling during the regenerative cycle), allowing us to observe how many and which teeth were being actively dissociated during the replacement process. Of the few oral tooth germs we observed (n=8 total observed from two separate fish) we found that stickleback oral replacement teeth formed at the labial side of each presumed predecessor tooth, and were not observed to be clearly diverging from a 1-for-1 replacement scheme (n=0/8 tooth germs) (Fig. 3A and B). In the pharynx, most stickleback replacement tooth germs appeared to be anchoring themselves solely in the place of a single presumed predecessor tooth, which was usually the tooth to which they showed the nearest epithelial association (Fig. 3C). However, in ∼25% of cases (17/67 tooth germs) a single replacement tooth was found to be abutting two erupted teeth that were both displaying interrupted mineralization and apparent concomitant osteoclast activity at their bases (Fig. 3D). We interpret these to be possible “1-for-2” tooth replacement events in the stickleback pharynx. These observations suggest that stickleback pharyngeal teeth deviate from a strict 1-for-1 mode of tooth replacement up to 25% of the time.

**Fig. 3.**
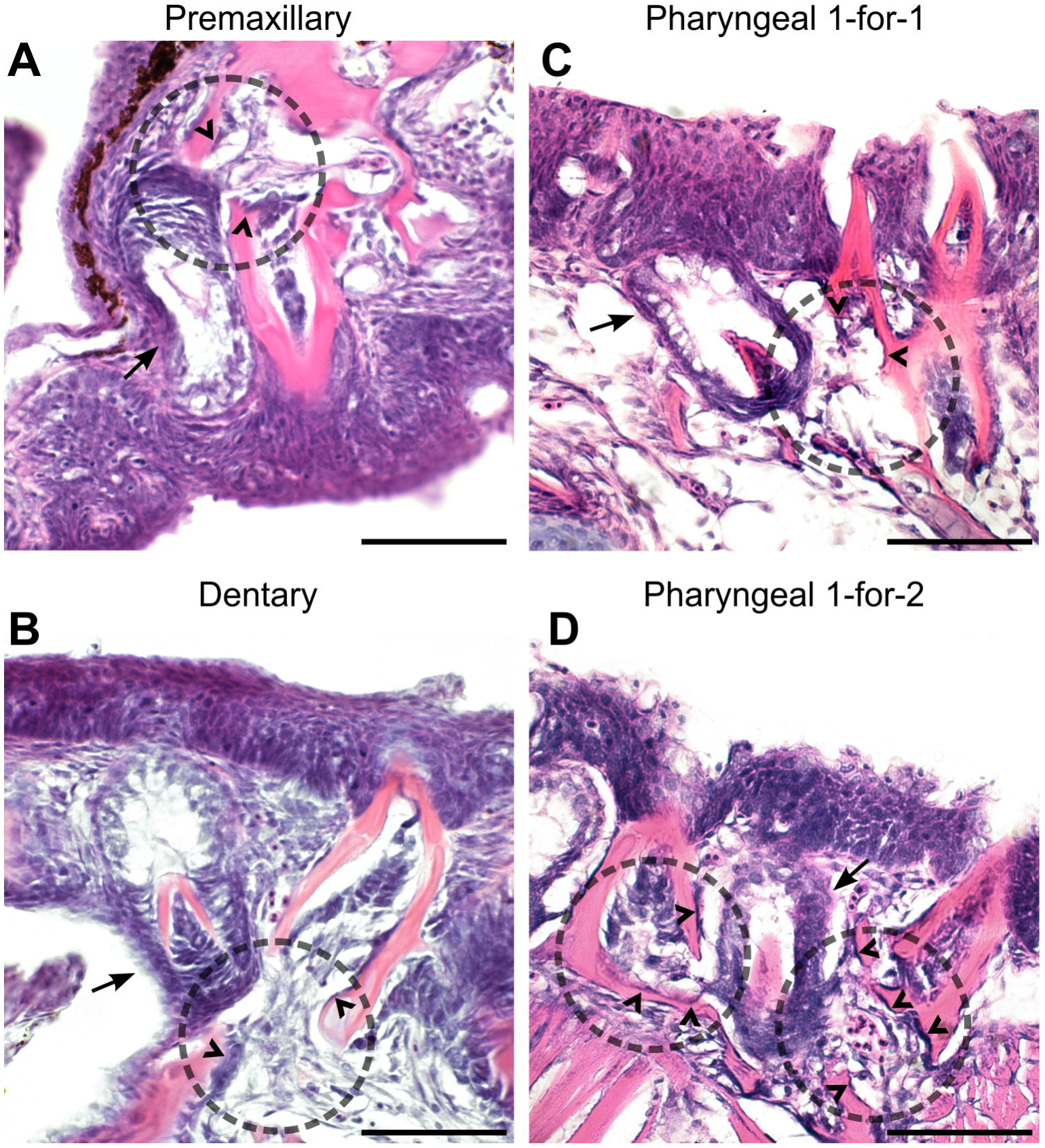
Oral and pharyngeal tooth replacement histology in stickleback. In all panels, dotted circles mark the general location of erupted tooth dislodgement adjacent to a presumed replacement tooth germ, arrows mark tooth germs, and carets indicate likely osteoclasts. All panels show sagittal (A, B) or transverse (C, D) sections and are oriented with the dorsal side facing upwards. **A, B** Oral tooth fields on the premaxilla and dentary bones were only observed as replacing via a 1-for-1 mechanism. **C** Pharyngeal replacement tooth germs were typically observed as dislodging only one other erupted pharyngeal tooth in 74.6% of cases (n=50/67, see Methods). **D** By contrast, some pharyngeal replacement tooth germs appeared to be dislodging two erupted teeth simultaneously in 25.4% of cases (n=17/67, see Methods). All scale bars = 20 μm.

### 2. Gene expression in stickleback and zebrafish tooth germs and progenitors

#### a. Zebrafish and sticklebacks sharply upregulate Wnt signaling in replacement tooth germs

The role of Wnt signaling in tooth morphogenesis and differentiation has been well-established in mouse and zebrafish [68–72], and indicted in multiple human tooth phenotypes [73–80]. Using *in situ* hybridizations on thin sections through zebrafish and stickleback tooth plates, we compared the expression of *Wnt10a* and *Lef1* in zebrafish and sticklebacks (Fig. 4A-D). We found highly conserved patterns of gene expression in tooth germs for both genes, with the expression of both *Wnt10a* and *Lef1* upregulated in the future inner dental epithelia (especially at the central epithelial cells of the placode, black arrows in Fig. 4A-D) and early condensing mesenchyme of bud stage tooth germs (white arrows in Fig. 4A-D).

**Fig. 4.**
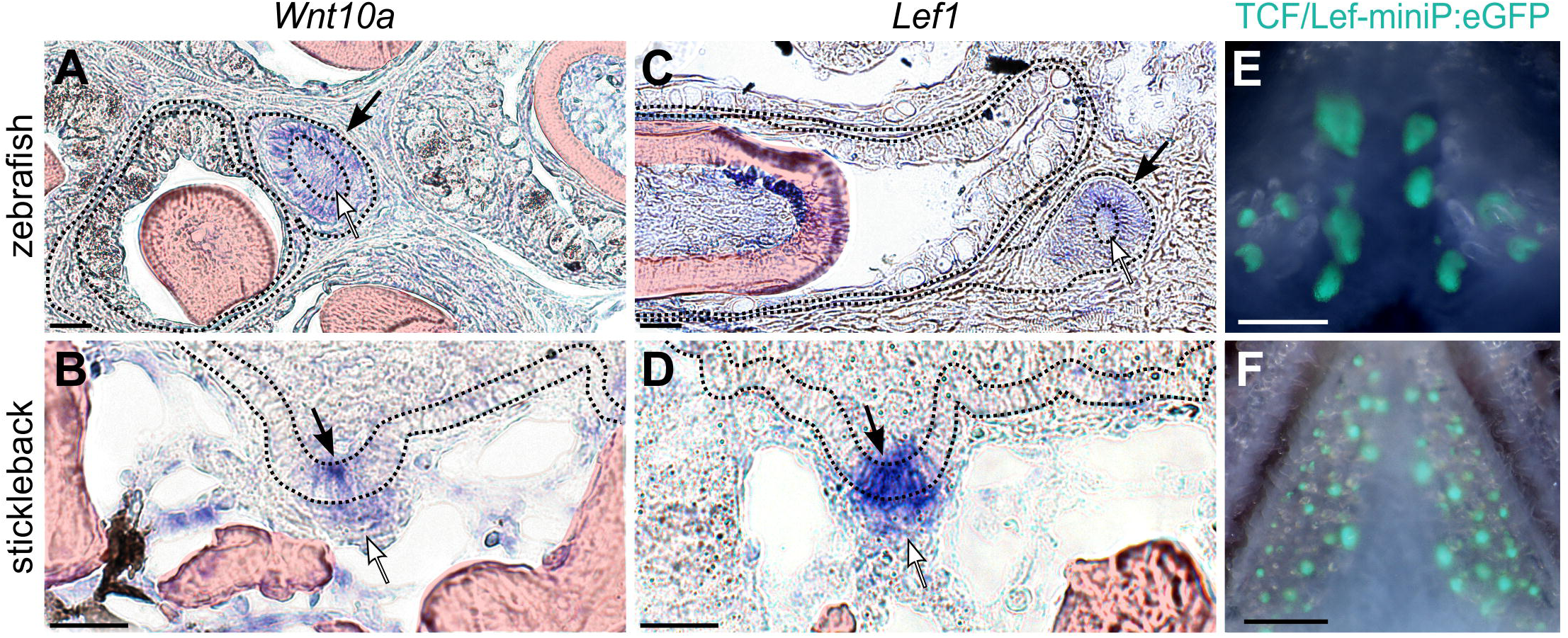
Wnt signaling in zebrafish and stickleback replacement tooth germs. Black arrows mark replacement tooth germ epithelium, white arrows mark replacement tooth germ mesenchyme. **A** *wnt10a* expression in a cap-stage zebrafish replacement tooth germ. **B** *Wnt10a* expression in a bud-stage stickleback replacement tooth germ. Note concentrated expression in the distalmost epithelium (arrow). **C** *lef1* expression in a cap-stage zebrafish replacement tooth germ. **D** *Lef1* expression in a bud-stage stickleback replacement tooth germ. **E, F** TCF/Lef synthetic Wnt activity reporter is active in adult replacement tooth germs. Scale bars in A-D = 20 μm; E and F = 500 μm.

One major output of Wnt signaling is upregulated transcriptional activity via β-catenin and TCF/Lef transcription factors [81]. A synthetic TCF/Lef reporter construct [82] was found to be expressed in a small number of cells in the young primary teeth of zebrafish, but expression was not detected in the first replacement tooth [72]. This result raised the possibility of a Wnt signaling disparity between primary teeth and replacement teeth. The construct used in these assays employed a destabilized enhanced GFP (dGFP) coding region, which works to add temporal resolution to the reporter construct output by increasing the degradation rate of eGFP. To address this potential primary vs. replacement tooth difference, we first compared the activity of the previously-tested *TCF/Lef-miniP:dGFP* reporter construct [82] in zebrafish and stickleback adults by generating novel transgenic lines in each species. We confirmed previous observations by Shim *et al*. (2019) that post-larval zebrafish teeth do not display detectable dGFP fluorescence (n=4/4 juvenile zebrafish). Similarly, we also did not detect dGFP by fluorescence microscopy in stickleback teeth at any stage. Given our *in situ* data for *Wnt10a* and *Lef1* expression in juvenile and adult fish (Fig. 4A-D), we suspected that downstream TCF/Lef transcriptional activity was also occurring in replacement tooth germs, but perhaps at a slightly reduced relative intensity and/or with faster turnover or signal dilution compared to the earliest zebrafish tooth germs. Thus, we re-stabilized the dGFP reporter cassette by removing the destabilization signal (see Methods) and again derived stable transgenic lines in zebrafish and sticklebacks. This change from dGFP to eGFP allowed the new *TCF/Lef-miniP*:*eGFP* cassette to be robustly detected in all developing young tooth germs in adult fish, especially tooth epithelium, and throughout the entire process of tooth morphogenesis (Fig. 3E and F). These data suggest that active Wnt signals are indeed transduced in replacement teeth of all ages in zebrafish and sticklebacks.

#### b. Expression in naïve epithelial tissues

As outlined above, sticklebacks lack a histologically-defined SDL (per [28]) in their pharyngeal tooth fields. Despite this, sticklebacks must still possess epithelial replacement tooth progenitors in the pharynx, given that pharyngeal replacement teeth form in part from the endoderm adjacent to erupted teeth (Fig. 1). Here we refer to these putative epithelial progenitors collectively as the “successional dental epithelium” (SDE). We use this general term to encompass both the cell types found in permanent or transient SDLs of other vertebrates, as well as the putative epithelial tooth progenitors of less-organized systems such as stickleback pharyngeal teeth.

Previous expression assays revealed that β-catenin, Plankoglobin, and E-Cadherin are expressed in the zebrafish SDL [83,84]. We aimed to bolster our understanding of gene expression in naïve dental epithelia. Using candidate genes from other known epithelial appendage precursor cell types (namely hair follicle stem cells), we performed *in situ* hybridization (ISH) on thin sections through zebrafish and stickleback pharyngeal tooth plates. To assess a variety of protein classes, we chose to assay orthologous genes in each fish species encoding three transcription factors (*Gli1, Nfatc1*, and *Pitx2*), three transmembrane receptors (*Bmpr1a, Lgr4*, and *Lgr6*), two secreted ligands (*Bmp6* and *Igfbp5a*), and a cell adhesion glycoprotein (*CD34*).

In zebrafish, we found orthologs of all nine genes (*bmp6, bmpr1aa, cd34, gli1, igfbp5a, nfatc1, lgr4, lgr6, pitx2*) to be expressed in the naïve, transiently-forming SDL during its earliest stages, when it consists only of a single epithelial bilayer (Fig. 5A). Other than in the dental epithelium of early tooth germs (presumably due to the maintenance of this genetic battery), the naïve SDL and its presumed daughter cells in young tooth germs were therefore the only observed pharyngeal cell types exhibiting expression of all nine of these genes. These data suggest a unique genetic signature of an SDL. One zebrafish co-ortholog we tested, *igfbp5b*, was not expressed in the SDL, and was instead expressed in a subset of dental mesenchyme (Fig. S3).

**Fig. 5.**
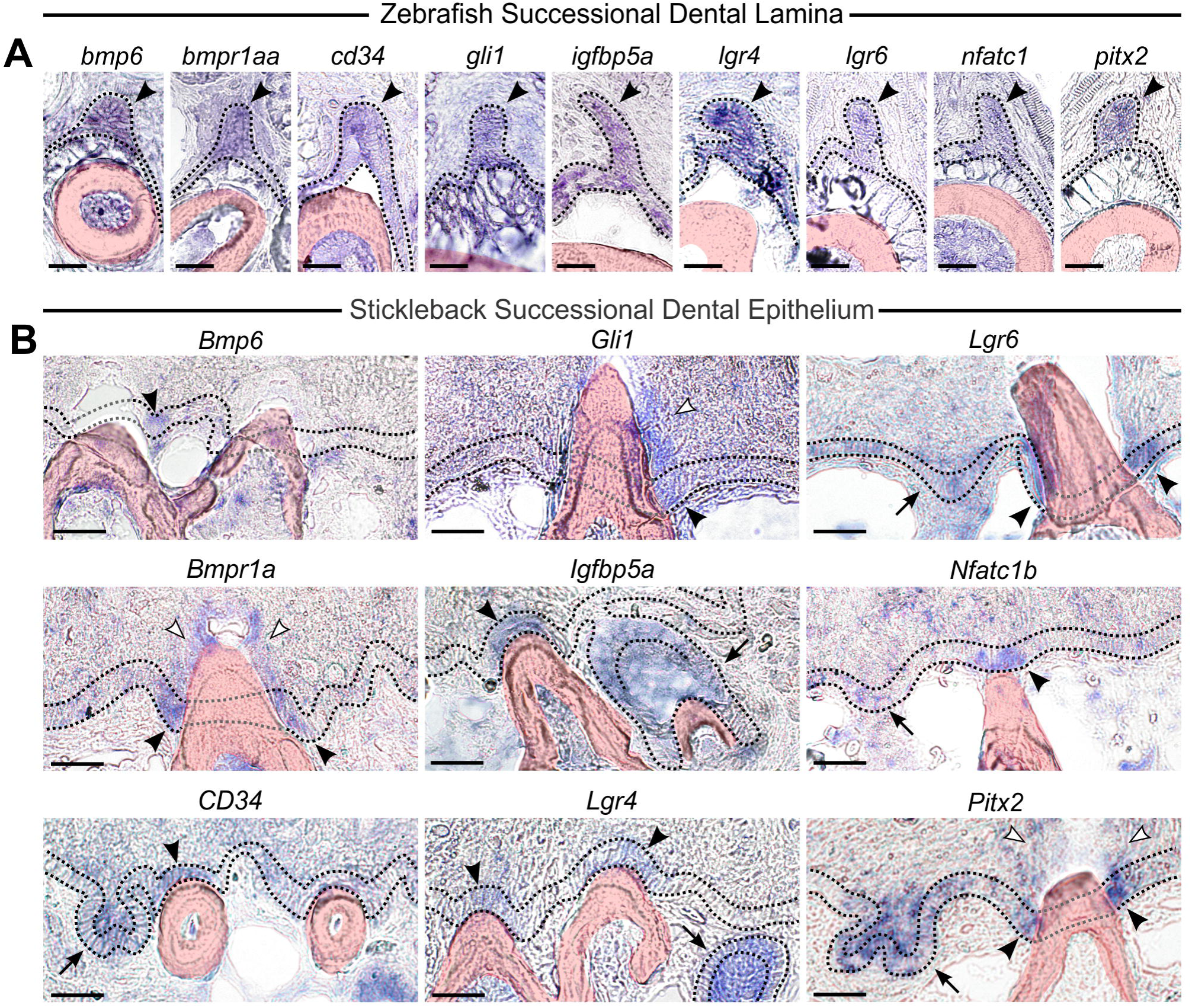
Successional Dental Epithelium (SDE) expression in zebrafish and stickleback. In all panels, a dotted line demarcates the basalmost pharyngeal epithelial layer, which is made grey when missing from a particular section (e.g. when it is interrupted by an erupted tooth). **A** Expression of *bmp6, bmpr1aa, cd34, gli1, igfbp5a, nfatc1, lgr4, lgr6, pitx2* were assayed by *in situ* hybridization (ISH) in zebrafish. Expression of all 9 genes was detected in the Successional Dental Lamina (SDL – black arrowheads). **B** Expression of *Bmp6, Bmpr1a, CD34, Gli1, Igfbp5a, Lgr4, Lgr6, Nfatc1b*, and *Pitx2* were assayed by ISH in stickleback. Expression of all 9 genes was detected in the proposed “Successional Dental Epithelium” (SDE) of stickleback. Black arrowheads mark the SDE. *Bmpr1a* and *Gli1* were additionally expressed in the superficial oral epithelium surrounding erupted teeth (white arrowheads). Tooth germs are indicated with a black arrow. All scale bars = 20 μm.

To test if the stickleback dentition deploys a genetic battery that is conserved with the zebrafish SDL anywhere in their pharyngeal tooth fields, we performed ISH on thin sections of stickleback pharyngeal tooth plates. We found that stickleback orthologs of all nine of the aforementioned genes (*Bmp6, Bmpr1a, CD34, Gli1, Igfbp5a, Lgr4, Lgr6, Nfatc1b*, and *Pitx2*) are expressed in the basalmost endodermal epithelium overlying pharyngeal tooth fields (Fig. 5B), specifically marking the “collar” of basalmost cells immediately juxtaposed to some, but not all erupted teeth, occasionally on flanking contralateral sides of a tooth (black arrowheads in Fig. 5B; see Fig. S2 for DAPI counterstain of nuclei of selected genes). Five of these nine genes’ expression patterns appeared locally restricted to the SDE (*Bmp6, Gli1, Igfbp5a, Lgr4*, and *Nfatc1b*), while the rest were found in more expansive stretches of the basalmost epithelium (*CD34, Bmpr1a, Lgr6*, and *Pitx2*). Three of these genes (*Bmpr1a, Gli1, and Pitx2*) had consistent detectable expression in the superficial pharyngeal epithelium surrounding erupted teeth (white arrowheads in Fig. 5). Importantly, as in zebrafish, this set of stickleback genes’ expression did not appear to delineate any other cell type in the pharynx, save early tooth germ epithelium (arrows in Fig. 5B), suggesting that this putative ‘SDE genetic battery’ is not common to other cell types or tissues in the pharynx. Notably, dental mesenchymal expression in differentiating and erupted zebrafish and stickleback odontoblasts was also observed for eight of these nine genes, to the exclusion of *Pitx2*, which was not detected in pharyngeal dental mesenchyme (Fig. S4). However, not all genes were expressed in all odontoblasts; in sticklebacks, *Bmp6* sparsely marked odontoblasts, while *Igfbp5a* and *Lgr4* only marked younger tooth germs (Fig. S4).

## Discussion

### 1. Successional Dental Epithelia

Across vertebrates, replacement tooth systems display variable arrangements, numbers of tooth replacement cycles, and levels of anatomical conspicuousness [9]. In some vertebrates, tooth replacement is anatomically well-defined and gives rise to many teeth per tooth position, while in other vertebrates, tooth replacement events are transient, ill-defined, or in some cases only give rise to a single tooth (like in humans and most other mammals that have only one wave of tooth replacement, a.k.a. diphyodonts). Here we asked if two vertebrates with histologically distinct tooth regeneration systems exhibit similar localizations of orthologous gene expression consistent with the presence of a conserved tooth epithelial progenitor cell type. We hypothesized that, despite the morphological differences found across regenerative systems, vertebrate teeth might deploy a homologous cell type responsible for generating the epithelial portion of a replacement tooth, but that this cell type may not always be found sequestered within a deep dental lamina. Here we show that zebrafish and sticklebacks, which are around 250 million years diverged [56], express a conserved battery of nine genes in their newly emerging replacement tooth epithelia. We propose this battery of gene expression labels a homologous, conserved dental cell type: the successional dental epithelium (SDE). Most of these nine genes have required functions in vertebrate dentitions, including *Bmp6, Bmpr1a, Igfbp5, Lgr4, Nfatc1*, and *Pitx2* [58,85–90], as well as other epithelial appendages (discussed below). Overall, these data suggest a common cell type (the SDE) underlies tooth regeneration in divergent tooth regeneration systems, regardless of the presence of a morphologically obvious lamina or SDL. More work in other vertebrates will reveal the level of gene expression domain conservation across the SDE of different vertebrate groups.

### 2. Is the lack of a distinct dental lamina plesiomorphic, or derived?

Micro-CT studies found that tooth plate fossils can retain signatures of past replacement events. In the fossil species *Andreolepis hedei*, a stem bony vertebrate (osteichthyan), tooth replacement did not appear to have occurred in a static 1-for-1 fashion, and likely lacked a permanent SDL [42]. These fossil data suggest that early modes of tooth replacement in ancient bony vertebrates are grossly similar to the condition in stickleback and other bony fishes such as bichir, medaka, and cod [18,44,67]. While some have concluded that such a regenerative system is likely secondarily derived [28], it remains possible that these species have retained an ancient, less orderly tooth regeneration system (i.e. it is plesiomorphic), and that the presence of a dental lamina could be convergent in certain vertebrate groups, as recently proposed [42].

By analyzing the histology and gene expression of other ray-finned fishes’ dentitions, it could be possible to test if the tooth regeneration systems like those found in sticklebacks and other fishes are more likely a plesiomorphic or derived character.

### 3. Tooth number evolution in sticklebacks

We found that stickleback replacement tooth germs initiate possible “1-for-2” replacement events up to ∼25% of the time. Our data thus leave open the possibility that adult tooth number variation in stickleback populations could be modulated by modifications to the replacement process, as previously posited [59]. Specifically, Ellis *et al*. (2015) found that marine and freshwater stickleback populations exhibited a consistent difference in adult (∼40 mm standard length) bilateral ventral tooth plate number, with an average difference of around 25 teeth, an approximately 50% increase in tooth number in derived freshwater fish relative to ancestral marine fish. In this same study, tooth germ number was quantified on ventral tooth plates, and found to have an average difference of ∼12 tooth germs in 40 mm adult fish, with freshwater fish from two different populations having more tooth germs than marine fish. As was previously addressed [59], these data suggest that the increased tooth numbers in freshwater stickleback populations relative to ancestral marine populations are likely not due to only differences in the appearance of new tooth germs; a difference in tooth shedding rate must also be at play. In line with these previous observations on inferred shedding rates, new histological data presented here suggest that the germ:shed ratio could be effectively doubled for some replacement events (1-for-1 vs 1-for-2) which conceivably could contribute to the overall tooth number differences seen between different populations. We hypothesize that up-or down-regulation of these “1-for-2” events may also contribute to these evolved tooth number differences in sticklebacks. Comparative studies on replacement events between multiple marine and freshwater stickleback populations could determine if “1-for-2” events differ in frequency between marine and freshwater fish, and if these differences could be related to genetic or developmental modifications to cell behavior or regeneration potential of the SDE.

4. Broader implications in epithelial appendage evolution and diversification Ideas surrounding the homology of disparate epithelial appendages (EAs), like teeth and hair, have been proposed as early as Darwin’s *On the Origin of Species* [91], and were thereafter bolstered by morphological comparisons and pleiotropic phenotypes affecting teeth, hair, and other epithelial organs [92–94]. Some consensus regarding the homology or shared nature of the placode stage has been established [54,55], while other work has indicated that larger-scale patterning mechanisms (e.g. concerted activity of Wnt, Eda, FGF, and SHH pathways) are at play across diverse EAs [48]. Distinguishing between uninterrupted ‘historical homology’ vs. deep homology [95,96] of EAs like teeth and hair remains difficult to assay using only extant vertebrates; fossil evidence of intermediate forms could feasibly provide links between modernly disparate EAs. Although it is not clear whether teeth and hair are related by strict-sense historical homology or cooption, teeth, hair, and other epithelial appendages exhibit widespread similarities in their early development and regeneration [20,53]. For example, in some rare instances, human dentitions form hair [97,98], and *Med1* mutant mouse dentitions form hair in the mouse incisor socket [99].

Here, we tested the hypothesis that two different tooth regeneration systems (zebrafish and stickleback) deploy similar patterns of gene expression (e.g. whether putative replacement epithelia have similar patterns of gene expression). Supporting this hypothesis, we found expression of a battery of nine candidate genes present in both the stickleback and zebrafish naïve dental epithelia, which we termed the SDE. Intriguingly, all nine of these genes are also expressed in mouse hair follicle stem cells, where most of these genes are known to play important roles in hair regeneration [62,63,100–107]. This similarity in gene expression suggests that hair and teeth may regenerate using a shared epithelial cell type, namely one that responds to Bmp and Wnt signals during the regenerative cycle, with Bmps suppressing and Wnts facilitating differentiation. This work builds on previous studies on stickleback gene expression levels in ventral tooth plates of 254 known hair follicle stem cell gene orthologs, where this gene set was found to be reduced in *Bmp6* mutants relative to wild type [58], and elevated in a high-toothed freshwater population relative to an ancestral low-toothed marine population [60]. Together, the expression profile similarities between hair follicle stem cells and dental tissues shown previously and in this manuscript will hopefully encourage further studies describing the shared and divergent genetic developmental mechanisms at play in these diverse organs.

One curious feature of fish tooth regeneration addressed previously is the apparent lack of slow-cycling label-retaining cells (LRCs) in the bichir and salmon oral dentitions, which do not employ an SDL [18,108]. Given that LRCs are present in other tooth regeneration systems [17,38,45,109] and in the hair follicle stem cell niche [110–114], these findings could represent a major difference between some fish dentitions and other epithelial appendages, including other fish teeth [45]. Further studies on LRCs in other epithelial appendages, including other fish species’ teeth, will help delineate how common and/or consequential these changes in cell behavior might be. However, regardless of whether LRCs or a true epithelial stem cell niche exists across all fish dentitions, further elucidation of the shared gene expression and cellular processes between the SDE, hair follicle stem cells, and other epithelial appendages is warranted and ongoing, including defining functional responses to Bmp and Wnt signaling inputs, and testing function of genes that mark the SDE and/or mesenchymal tooth precursors during tooth regeneration.

## Methods

### 1. Animal husbandry

Animals were raised following UC Berkeley IACUC protocol AUP-2015-01-7117. Zebrafish husbandry adhered to standard methods [115]. Sticklebacks were raised in 110L aquaria in brackish water (3.5g/L Instant Ocean salt, 0.0217g/L sodium bicarbonate) at 17-18° C in 8 hours of light per day. Fish were fed live artemia as young fry, live artemia and frozen daphnia as young juveniles (1-2 cm standard length), frozen daphnia and frozen bloodworms as older juveniles (2-3 cm standard length), and frozen bloodworms and *Mysis* shrimp as sub-adults and adults (3+ cm standard length).

### 2. Generation of *TCF/Lef-miniP* stable transgenic lines

The *TCF/Lef-miniP:dGFP* plasmid was a gift from Tohru Ishitani [82]. To ‘restabilize’ GFP, we removed the destabilization signal by digesting the previously published plasmid with HindIII and ClaI, and filling this lesion with a short double-stranded piece of DNA comprised of two synthesized (Integrated DNA Technologies), phosphorylated, and annealed oligonucleotides. Forward oligo: 5’ AGCTTTAAAGCGACAT 3’, reverse oligo: 5’ CGATGTCGCTTTAA 3’. This change to the plasmid removed the majority of the destabilization signal, leaving just 2 extra amino acids (Lys, Leu) before a stop codon on the C terminus of eGFP. Zebrafish and stickleback transgenic lines were generated using both the unmodified and modified versions of the *Tcf/Lef-miniP* constructs according to standard methods in each species, using Tol2-driven transgenesis [116] by injecting zygotes at the one-cell stage [117,118].

### 3. Alizarin Red staining

Alizarin Red was used to stain tooth plates as described previously [119]. In brief, whole fish were fixed in 10% neutrally buffered formalin overnight at room temperature, and washed with tap water or 1x phosphate-buffered saline with 0.1% Tween-20 (PBST) for 1 hour at room temperature. Samples were then washed in a 0.008% Alizarin Red S solution in 1% KOH for 24+ hours. Once adequately stained, samples were transferred to 1% KOH for 1-3 days to continue clearing. Once cleared, tooth plates were dissected, further cleared in 50% glycerol for 1-5 days, flat-mounted, and imaged on a Leica DM 2500 compound microscope.

### 4. Sectioning

Stickleback sub-adults (∼2 cm) derived from marine (Rabbit Slough [RABS]) and/or freshwater (Cerrito Creek [CERC] or Fish Trap Creek [FTC]) populations, and AB strain zebrafish adults or sub-adults (1.5-2 mm) were euthanized, decapitated, and fixed overnight in 4% formaldehyde (Sigma P6148) in 1x phosphate-buffered saline (PBS) at 4° C with heavy agitation, washed 3x 20 min with PBST on a nutator, then decalcified for 5-7 days (sticklebacks) or 3-5 days (zebrafish) in 20% ethylenediaminetetraacetic acid (EDTA, pH 8.0) at room temperature on a nutator. Once decalcified, specimens were again washed 3x 10 min in PBST, then stepped into 100% EtOH via 15-60 min washes in 30, 50, 70, and 100% EtOH in RNAse free H_2_O. Samples were sometimes washed into MeOH and stored at this stage for up to 12 weeks. Samples were then washed for 1 hr+ in 50/50 EtOH/Hemo De at room temperature, then 1 hr+ in 100% Hemo De at room temperature, then 1 hr+ in 50/50 Hemo De/paraffin (Paraplast x-tra, Fisher) at 65° C, then rinsed and washed overnight at 65° C in 100% paraffin. Fish heads were embedded in plastic molds with 100% paraffin (heated to 65° C), mounted, sectioned on the sagittal (stickleback), off-sagittal (∼30 degree tilt on the coronal axis; zebrafish), coronal (both species) or transverse (stickleback) planes with a Microm HM 340 E microtome. Sections were captured on Superfrost Plus slides, sometimes stored for up to 3 weeks at room temperature prior to analysis. To prepare slides for histological staining or ISH, slides were de-parafinnized (5 min incubation at 65° C, let cool, submerge for 5 then 10 min in 100% Hemo De, 5 min 100% EtOH, 10 min 80% EtOH/H2O, 10 min 100% H_2_O).

### 5. Hematoxylin and eosin (H&E) staining on sections

H&E staining was performed on sectioned tooth material using the following series of washes: Hematoxylin solution (VWR) for 3 min, tap water for 2x 20 seconds, bluing solution (0.1% Sodium bicarbonate) for 2 min, tap water for 20 seconds, acid alcohol (0.32% HCl in 95% EtOH) for 20 seconds, tap water for 20 seconds, eosin solution (VWR) for 30 seconds, 95% EtOH for 2x 2 min, 100% EtOH for 2x 2 min, then Hemo-De for 2x 5 min. Slides were then immediately coverslipped with Permount (Fisher), allowed to dry overnight, and imaged on a Leica DM2500 compound microscope.

### 6. *In situ* hybridizations on Sections

All in situ hybridization (ISH) steps were performed on freshwater (CERC) stickleback, or AB strain zebrafish sections. The following steps were performed in LockMailer microscope slide jars (Sigma-Aldrich) in a volume of 9-11 mLs unless stated otherwise. To begin the *in situ* process, slides were washed for 5 min in PBST, 5 min in proteinase K solution (15 μg/mL in PBST), rinsed briefly with PBST, then re-fixed for 20 min at room temperature in 4% formaldehyde in PBS. Fixative was then washed out with one PBST rinse and 2x 10 min PBST washes before pre-hybridization. Slides were washed 2x 5 min at room temperature in hybridization buffer, followed a long incubation in hybridization buffer (no probe, “pre-hyb” step) for 1-4 hours at 67° C in a rotating hybridization oven (hybridization buffer is 50% formamide, 5x SSC, 0.1% Tween, 5 mg/mL CHAPS, 1 mg/mL yeast RNA, 0.1 mg/mL heparin, pH 6.0 with citric acid). Riboprobes were generated essentially as described previously [64] against gene orthologs of interest (see Supp. Table 1). Riboprobe templates were created either by PCR cloning gene fragments from cDNA or genomic DNA, or by synthesizing whole or partial inferred coding sequences (Gene Universal, Delaware, USA). See Table S1 for a complete list of transcript accession numbers and sequences corresponding to exact riboprobe templates used in this study. Stickleback *Bmp6* and *Pitx2* riboprobes were previously published [57]. The zebrafish *pitx2* riboprobe was previously published [120]. Zebrafish and stickleback *CD34* orthologs were identified using the Genomicus synteny browser v100.01 (www.genomicus.biologie.ens.fr), which supported orthology of mammalian *CD34* to the fish genes assayed here (ENSGACG00000011016 and ENSDARG00000095268). The translated products of these gene appear to code for proteins containing a partially recognizable CD34 domain (pfam06365). Riboprobes were synthesized with digoxygenin-labeled UTP and added at a concentration of ∼100-500 ng/mL in 10 mL of hybridization buffer and agitated overnight in a rotating hybridization oven at 67° C. Riboprobes in hybridization solution were stored at -80° C and reused in some cases. The following day, six pre-heated hybridization washes at 67° C in a rotating hybridization oven were performed for 20–90 min each, totaling 5-6 hours of total “hyb wash” time (hyb wash is the same recipe as hyb buffer, excluding CHAPS, RNA, and heparin). Slides were then rinsed and washed in pre-heated Maleic acid buffer with Tween (MABT) at 67° C for 20 min, then washed in pre-heated MABT for 20 minutes at room temperature (to allow for slow cooling). Slides were then removed from slide jars, placed in a humidor (a sealed plastic container with wet paper towels for moisture and pieces of plastic to raise slides form the bottom), and blocked with 70-100 μl (enough to cover all tissue on slides) of 2% Boheringer Blocking Reagent (BBR), covered with parafilm for one to three hours at room temperature. Following the block step, block was poured off each slide, and anti-Digoxygenin Alkaline Phosphatase conjugated antibody (Roche SKU 11093274910) was added at a concentration of 1:2000 in 2% BBR, again using enough to submerge each piece of tissue on each slide (70-100 μl) beneath parafilm and incubated at 4° C overnight, or 3h at room temperature in the dark. Importantly, tissue was not allowed to dry out beneath the parafilm during the block or antibody steps above. The following day, we performed one MABT rinse and 5x 20-50 min MABT washes (back into slide jars) over the course of 3-4 hours, agitated at room temperature, to wash out residual antibody, usually followed with an overnight MABT wash unagitated at 4° C. To begin the coloration process, slides were changed into NTMT (0.1M Tris pH 9.5, 0.05M MgCl_2_, 0.1M NaCl, 0.1% Tween) via 3x 5-10 min washes before removing the final NTMT wash and replacing it with 10 mL of coloration solution (NTMT with 25 μg/mL Nitro blue tetrazolium chloride [NBT] and 175 μg/mL 5-bromo 4-chloro 3-indolyl phosphate [BCIP]). Signal development was carried out for 2-30 hours to visualize mRNA localization. Once adequately developed, slides were rinsed, then washed for 10 min in PBST, fixed in 4% formaldehyde in PBS for 1-5 days at 4° C. Slides were then washed 3x 5 min in PBST. At this point, some select slides were counterstained with 0.1 μg/mL DAPI in 1x PBST for 5+ min at room temperature (the same 10 mLs of solution were used to stain all sections shown in this paper, stored at 4° C). To prepare slides for imaging, they were rinsed then washed 3x 5+ min with deionized H_2_0, coverslipped with deionized H_2_0, and imaged on a Leica DM2500 microscope.

### 7. Quantification of 1-for-1 vs 1-for-2 tooth replacement in sticklebacks

To address the possibility of 1-for-2 tooth replacement in sticklebacks, we H&E stained serial sections (described above) of nine sticklebacks of various sizes (20-40 mm) from various backgrounds (5x CERC, 1x FTC, 2x RABS, 1x marine/freshwater hybrid). Across these preparations, we identified a total of 67 pharyngeal tooth germs for which we had complete histological profiles at the mid-to late-bell stage, that were also present in the middle of the tooth plate (i.e. surrounded by erupted teeth), which represent presumed or potential replacement teeth. Of these 67 tooth germs, 17 (25.4%) were clearly observed to be abutting and connected to bone remodeling events at the bases of two erupted teeth. These associations also appeared to contain an abundance of likely osteoclasts surrounding the disrupted bone in the erupted teeth (carets in Fig. 4), which we interpret as evidence of possible shedding events (or at the very least, loosening events) in-progress at the time of tissue preparation. We similarly located all oral tooth germs present on these slides, though the oral tooth fields were only visible on two sectioned specimens. We did not observe oral tooth germs as diverging from a 1-for-1 replacement scheme (n=0/8).

## Acknowledgements

We thank Andrew Glazer, Priscilla Erickson, and Justin Roncaioli for generating the *Lef1, Igfbp5a*, and *Lgr6* riboprobe templates, respectively, James Hart for assistance and advice on stickleback expression data, Amy Shyer and Richard Harland for microtome assistance and access, and Sophie Archambeault, Alyssa Bormann, and Mark Stepaniak for valuable feedback on the manuscript.

## Declarations

### Authors’ Contributions

TAS and CTM conceived the study. TAS designed the experiments. TAS, SBS, and EJM performed the experiments and collected the data. TAS prepared the figures. TAS and CTM wrote the manuscript. All authors provided input on the contents of the final manuscript.

### Availability of Data and Materials

The datasets used and analyzed during the current study are available from the first author on reasonable request.

### Ethics approval and consent to participate

Animals were raised following UC Berkeley IACUC protocol AUP-2015-01-7117.

### Consent for Publication

Not applicable

## Supplementary figure legends

**Supp. Fig. 1**. Histological plane orientation and description in sticklebacks. Illustrations at top depict a tooth (red cone) and indicate section planes with dotted rectangles, which are pictured below for each respective plane of section (labeled at top). These planes of section roughly translate to whole-animal anatomical planes. Left and middle illustrations show the sagittal and/or transverse plane (because stickleback teeth are mostly conical, these two planes of section appear generally similar in section). On these planes of section, teeth often appear off their medial axis (left vs center illustration, see labels). Rightmost illustration depicts the coronal plane, where teeth appear as rings of bone. Bottom images are of H&E-stained sections representing each illustrated plane of section above. Left bottom image shows a pharyngeal tooth on the transverse plane. Center bottom image shows an oral tooth on the sagittal plane. Right bottom image shows a pharyngeal tooth sectioned on the coronal plane. An erupted tooth (“et”) is indicated in the center image of an off-medial tooth (suggested with a red dotted line) sectioned on the sagittal plane. In all H&E images, tooth germs are indicated with an arrow, and the SDE is indicated with an arrowhead. All scale bars = 50 μm.

**Supp. Fig. 2**. DAPI counterstaining *in situ* hybridized sections aids in distinguishing epithelium from mesenchyme. Three gene expression patterns are shown as examples, labeled at top of columns in figure. Each column shows the same section corresponding to each gene, top two images are in brightfield, bottom two show DAPI fluorescence. 1^st^ row of brightfield images are shown without any markup. 2^nd^ row of brightfield images outline the basalmost epithelium with black dotted lines where possible (gray segments suggest borders out of the plane of section); bone is false-colored red. Expression domains in tooth germs are indicated with black arrows. Expression in the SDE is indicated with black arrowheads. Superficial epithelium with *Gli1* expression adjacent to erupted teeth is indicated with white arrowheads. 3^rd^ row of images shows the DAPI counterstain of each brightfield image above it, with the same epithelium outlines overlain (white dotted lines). 4^th^ row shows DAPI images without any markup. Scale bar = 50 μm and applies to all panels.

**Supp. Fig. 3**. Zebrafish *igfbp5b* expression in deep tooth mesenchyme. Top row shows brightfield images, bottom row shows the corresponding DAPI counterstains to each brightfield image above. The SDL of zebrafish did not show *igfbp5b* expression (gray arrowhead). Expression was detected in a subset of the deep dental mesenchyme (white arrows). Scale bar = 50 μm and applies to all panels.

**Supp. Fig. 4**. Expression of *Bmp6, Bmpr1a, CD34, Gli1, Igfbp5a, Lgr4, Lgr6, Nfatc1*, and *Pitx2*, highlighting mesenchymal expression. All of these genes were found to mark the SDE, except for *Pitx2*, and are expressed in some portion of dental mesenchyme (white arrows). Species and gene assayed are indicated in the figure. All scale bars = 50 μm.

