## Supplementary figures and images for "Distinct tooth regeneration systems deploy a conserved battery of genes"

### Fig. S1

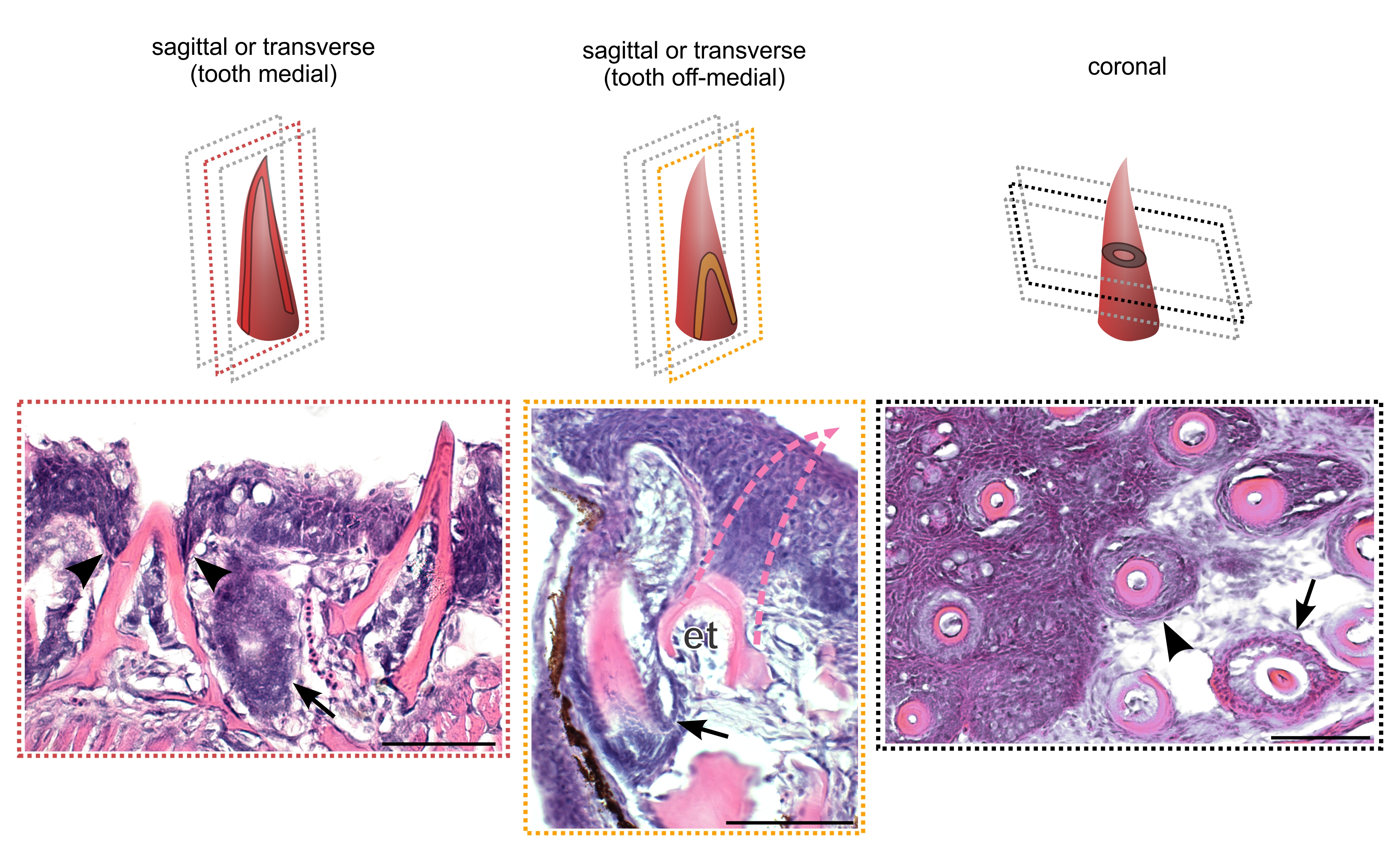

### Fig. S2

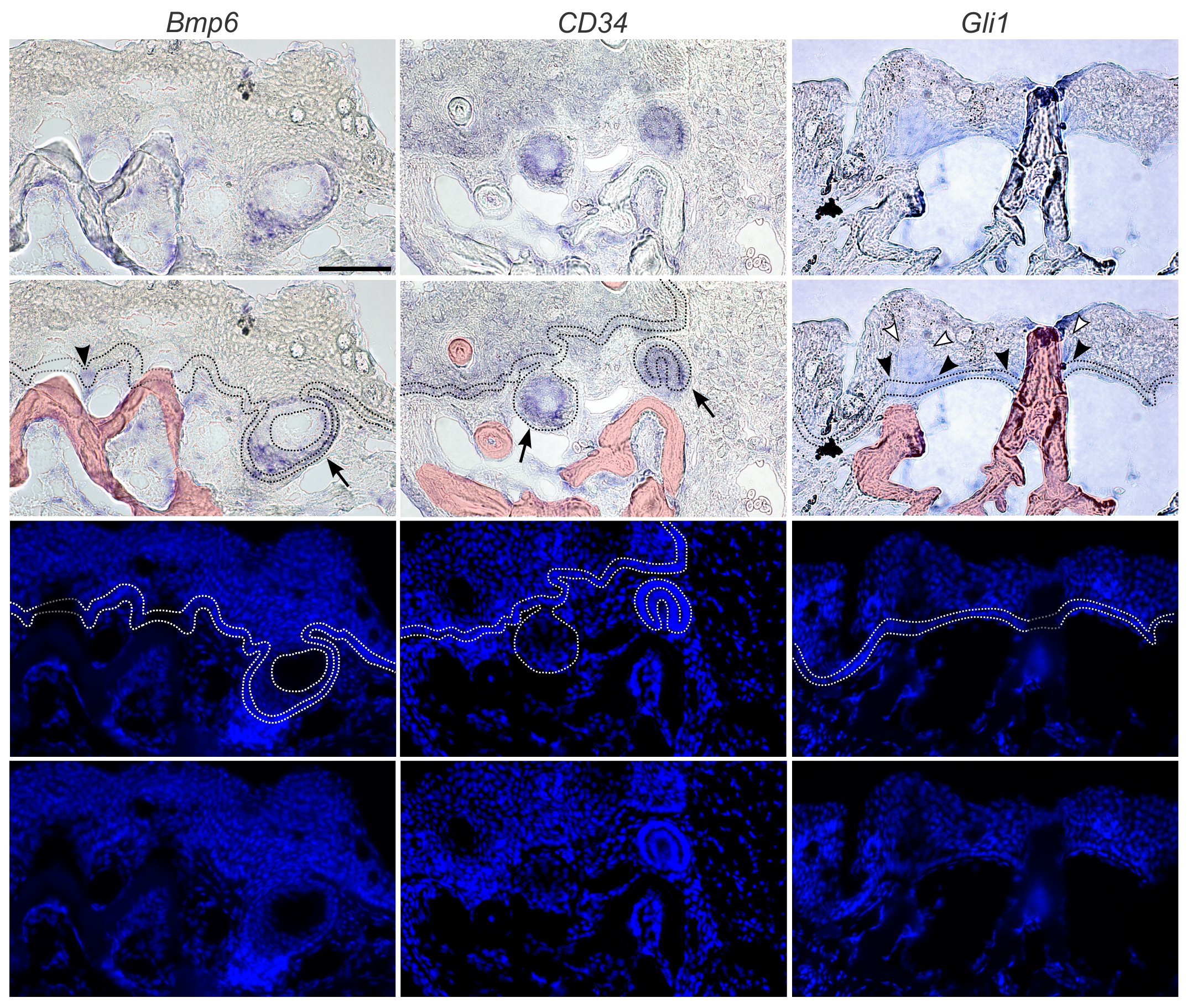

### Fig. S3

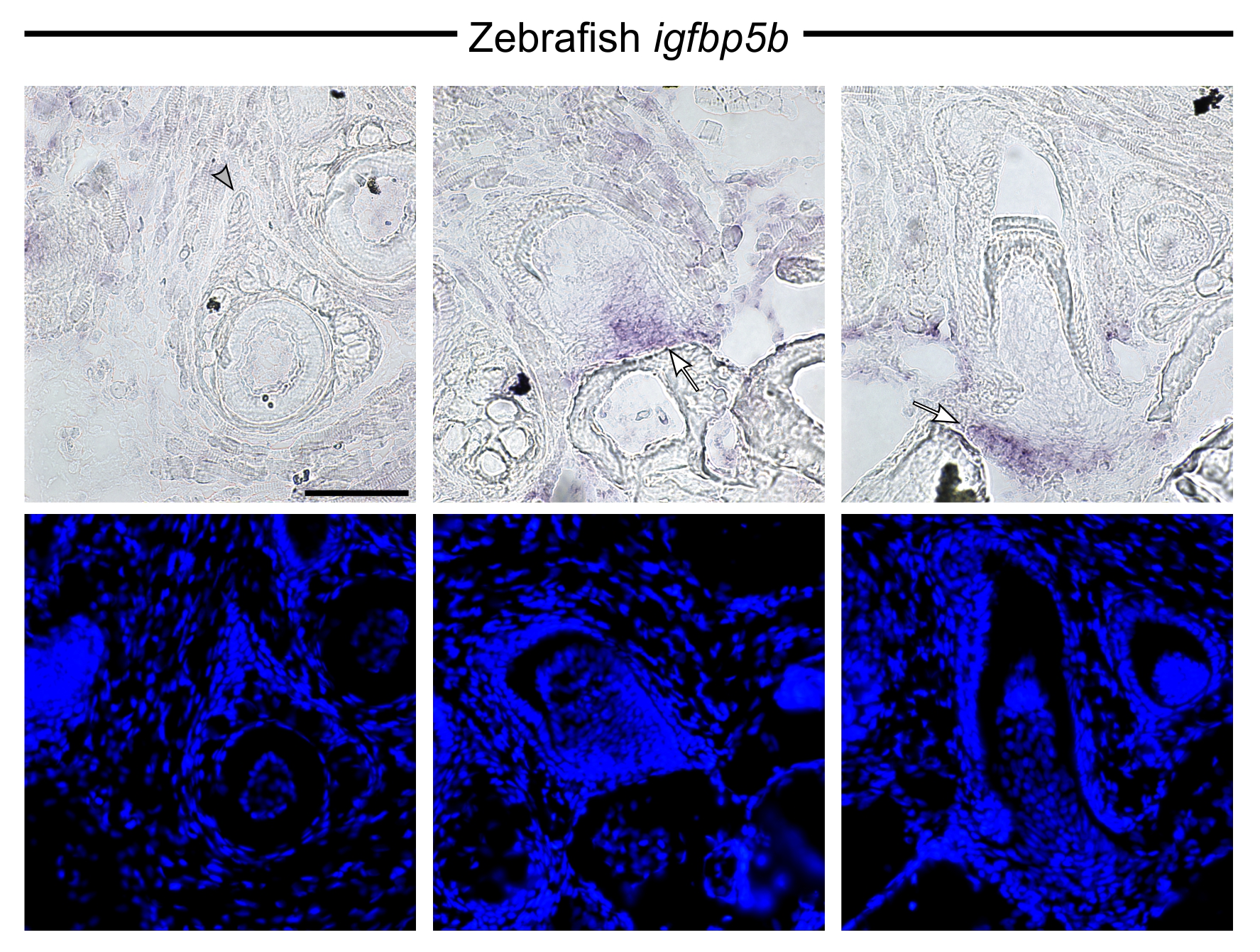

### Fig. S4

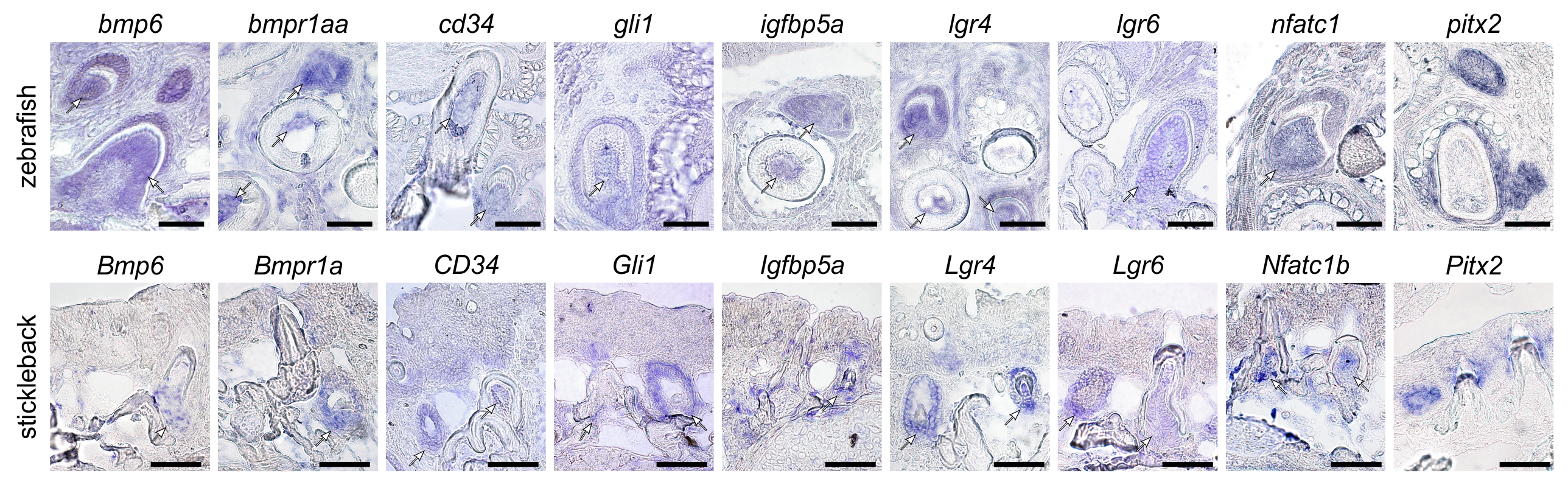
